# Modelling the willingness to pay for honey: the role of sensory characteristics and EEG neurometrics

**DOI:** 10.1101/2025.01.30.635681

**Authors:** Julia Zaripova, Daria Semenova, Anna Provorova, Sofya Kulikova

## Abstract

It is currently unclear which honey attributes are most important for consumers and, more importantly, from an economic perspective, for which of them consumers are willing to pay for. The aim of this study was to examine the role of sensory attributes in consumers’ willingness to pay for honey. The study was performed in two steps. First, we ran a behavioural experiment in which 25 participants were asked to blindly taste and evaluate 21 sensory attributes of 7 different flower honey samples. Based on the obtained data, we conducted a factor analysis and aggregated all sensory attributes into 6 factors, which were used in the second part of the study during a neuromarketing experiment. The neuromarketing experiment involved another 25 respondents who also blindly tasted honey, evaluated its 6 sensory characteristics (corresponding to the 6 above-mentioned factors), as well as their willingness to pay, the perceived quality and taste of the samples. While the respondents were evaluating honey samples, their brain activity was recorded using an electroencephalograph (EEG). Based on the obtained data, the willingness to pay was modeled using a linear regression, which has demonstrated that consumers’ willingness to pay for honey is significantly influenced by the taste of honey, the perceived quality indicated by various sensory characteristics, the male gender, and the brain activity in the frontal areas.

## 1. Introduction

Honey and other apiculture products are the fairly popular categories, regularly found in the food basket of modern consumers. Researchers note a growing trend in honey consumption, which is caused by the increasing interest in healthy food of organic origin. Honey is positioned as a healthy food product due to its composition. It may be used as a substitute for refined sugar in hot and cold beverages, as a traditional ingredient in cooking for cakes, desserts, sauces, sandwiches, and cheese (Kowalczuk, 2023). The antioxidants, phenols, and flavonoids contained in honey have favourable effects on mental health, improving memory and helping to cope with stress (Zamri et al., 2023). The motives for consuming honey are rather diverse. The product is widely discussed as an adjuvant folk remedy to treat upper respiratory tract diseases and to reduce body temperature, to heal wounds, to support the immune system, to decrease cholesterol levels and to prevent some forms of cancer (Kumar, 2010). Honey is often used for cosmetic purposes to moisturise, purify and improve skin elasticity, to relieve irritation and to slow down the ageing processes. Modern consumers also actively mention the nutritional properties of honey (Grontkowska & Grzyb, 2019). In this paper, we primarily discuss honey as a food product with the goal to identify its key sensory characteristics and their contribution to the perceived quality and willingness to pay (WTP).

Existing studies on consumer WTP for food products focus on identifying factors related to perceived taste (Raimondo et al., 2024) or quality (Wu et al., 2015). While some researchers have explored the impact of packaging on honey’s perceived value, few studies have delved into the assessment of sensory characteristics, which are likely to play a crucial role in influencing repeated purchases after the initial tasting experience. Therefore, one of our objectives was to investigate how sensory characteristics of honey influence consumers’ WTP and how they contribute to the perceived taste and quality. By examining these relationships, we seek to shed light on the factors that drive consumer preferences and purchasing decisions in the context of honey products. In particular, we were interested in finding answers to the following questions:

- Which honey characteristics positively influence the perceived product quality?
- How do the perceived taste and quality increase consumer WTP?

Apart from these questions, we also wanted to investigate how honey perception and the WTP for it change with age and gender and whether individual purchasing decisions are reflected by some cognitive processes that could be evaluated using EEG-based neurometrics.

The following text is organized as follows. First, in Section 2.1, we discuss factors that may influence the perception of honey. Then, in Section 2.2 we make some notes on using EEG for investigating WTP. Section 3 describes the methodology of the present work: Section 3.1 gives details on the behavioral part of the study, while Section 3.2 describes the neuromarketing experiments. The results are presented in Section 4 and they are discussed in Section 5.

## 2. Theoretical background

### 2.1. Factors influencing the honey purchase decision

Honey is a “complex” food product, so the decision to purchase it is influenced by a wide range of factors (Sparacino, 2022). Previously, three key variables influencing consumers’ WTP have been identified, namely distribution channel, packaging and price. Although most studies were focused on the effects of externalised product characteristics (price, packaging material, point of sale, location of honey producer), consumers decide to re-purchase honey after tasting it and evaluating internalised characteristics (taste, aroma, texture, etc.), making it important to investigate how sensory characteristics influence the perceived quality of honey.

There is a large variety of sensory characteristics used to describe the flavour “portrait” of honey. Producers predominantly operationalise most of them for expert evaluation of the product quality. In this study, we were interested in product properties that can be easily evaluated directly by common consumers. In order to increase their WTP, it is necessary to find out the terms they use to describe different honey samples.

The most frequently encountered sensory characteristics of honey across the literature include colour and odour intensity, texture (liquid, crystalline), florality, fruitiness, degree of waxy, chemistry, fermentability, bitterness, astringency, sourness and acidity, mouthfeel, and degree of aftertaste (Hunter, 2021). Let us describe these factors below in more detail.

#### Texture

Commonly, honey texture refers to the perceived degree of crystallisation. According to the type of texture, honey may be liquid, creamy or crystallised. Most often, customers prefer liquid honey: it seems to be sweeter and more delicate. As the honey crystallises, its sweetness decreases, and the firmness and graininess increase (Piana et al., 2013). It is believed that consumers trust liquid honey because they perceive it to be fresher (Šedík, 2023). On the other hand, in some studies, respondents preferred creamy honey to liquid (Khaoula, 2019).

#### Aroma

Counterfeit usually has a less intense aroma, so consumers may focus on this indicator in a honey selection (Šedík, 2018).

#### Flavour

The sweetness of honey can be identified as the most attractive characteristic for consumers. Most of the other sensory characteristics, according to the work of Hunter (2021), have a negative impact on the attractiveness of honey. The same study points to a link between sweetness and the perceived amount of sugar in the product, which, in its turn, suggests the participants perceived honey as a hedonic product.

The set of sensory attributes used to describe honey varies across countries, showing some cultural specificity. For example, a term “Jaggery-like”, which stands for aroma associated with unrefined brown sugar made from palm sap, may be found in Indian studies (Anupama, 2003), while “Animal-like” characteristic, which reflects a honeydew flavour, may be reported in Baltic regions (Kivima, 2021). Thus, when describing honey flavour profile, it is also important to take into account regional specificity. Therefore, in our behavioural experiment, we considered an additional characteristic familiar to local honey consumers called “peppercorn” (the tickling sensation in the throat after the honey tasting). In total, 21 sensory characteristics were identified as potentially relevant and included into the initial list to describe the taste of honey (see Section 3.1.3 for the full list of characteristics).

Previous works argue that honey perception is influenced not only by product-related characteristics but also by respondents’ personal attributes such as gender and age (Sparacino et al., 2022). The study by Zeng et al. (2023) indicated that men buy honey significantly less frequently compared to women, and as a consequence, their WTP may differ from those of the female audience. In addition, older consumers are more likely to purchase honey due to the fact that it is considered as a traditional healthy treat in their minds (Zeng et al., 2023).

### 2.2. Using electroencephalography for evaluation of consumers’ perception of food products

For better understanding the factors of consumers’ choice, and identifying the drivers of positive purchasing behaviour, both academic and business communities turn towards implementation of the qualitative and quantitative marketing research methods. It is important to note that traditional and familiar tools — questionnaires, interviews, focus groups, and observations — are designed to study the conscious component of the decision-making process, while most of its stages are governed by subconscious processes (Agarwal, Dutta, 2015).

In the early 2000s, an alternative approach to study consumer behaviour appeared, named neuromarketing. Neuromarketing is a new interdisciplinary field at the intersection of marketing, psychology and neurobiology (Plassmann et al, 2012). Neuromarketing aims at studying the neuropsychological and emotional mechanisms that explain the behaviour of individuals, in particular, the purchase decision-making process (Alvino et al, 2020).

The choice of food products seems to be a complex process, which is currently defined as under-investigated (Stasi et al., 2018). The list of classical parameters of product audience segmentation, often determined directly by the respondents of the study (socio-demographic characteristics, stated preferences, etc.), does not include the influence of some attitudes and emotional reactions to the product on consumer choice. Thus, using the neuromarketing tools may help to reduce misinterpretation of the respondents’ choices.

In this study, the neuromarketing metrics (or neurometrics) were based on electroencephalography (EEG) data. EEG presents a method that allows the recording and evaluation of the electrical activity of the cerebral cortex that is generated by neuronal cells (neurons) in response to various stimuli and during the decision-making process. The method is the method of choice for the majority of neuromarketing studies thanks to its important advantages: non-invasiveness, ease of performing the diagnostic method, the ability to conduct experiments repeatedly, high temporal resolution, high sensitivity to changes in bioelectrical activity of the brain. It has been shown that EEG can effectively capture consumers’ physiological and emotional reactions to food products when participants are exposed to different images, sounds, taste, and odours, and the recorded data may help to explain the peculiarities of product perception (Songsamoe et al., 2019).

However, the number of stimuli to be tested and the duration of the experiment should not be too long to avoid fatigue of the participants and signal deterioration due to drying of the electrodes. Given the above fact, we decided to conduct our study in two stages: in the first stage, one group of respondents tasted and evaluated 21 characteristics of honey. Next, we processed this data using factor analysis and aggregated the 21 sensory characteristics into larger components. We used these components at the second stage of the study - in a series of neuromarketing experiments, during which another group of respondents also tasted honey blindly, evaluated their WTP, the perceived quality and taste, as well as the aggregated sensory characteristics. Each stage of the study is described below in more detail.

## 3. Materials and method

### 3.1. Behavioral experiment

#### 3.1.1. Participants

Twenty-five participants (16 female / 9 male, aged 18 to 65 y.o) were enrolled in the first stage of the study. The respondents were recruited according to the following criteria: participants had to be over 18 years old (to exclude the effect of parental budgeting), without allergies to honey and apiculture products or other contraindications for honey degustation. Each participant had experience of buying honey in a supermarket at least once. No restrictions regarding gender, education level or profession were applied. Respondents were recruited via snowball sampling through popular social media. All the respondents gave their written informed consent to participate in the study, successfully completed the experiment and received a gratification of 250 monetary units.

#### 3.1.2. Experimental design and stimulus materials

The behavioural part of the study consisted of two distinctive stages and lasted 1 hour and 15 minutes in total.

##### Preparatory stage

Before the beginning of the session, all participants were instructed: the moderators explained the purpose and the content of the experiment, informed about potential allergic reactions to honey. At this stage, participants filled out paper-based questionnaires about their socio-demographic characteristics, traditional behaviour in terms of buying and consuming honey, and the current level of satiety.

##### Degustation

During this stage, respondents were asked to taste seven samples of honey and fill out a questionnaire form to evaluate each sample. The time for tasting the samples was not limited. Participants could also correct their answers during the tasting. Water was used as palate cleansing material in between the samples. The amount of water was not limited. The amount of honey of each sample was limited to 50 g, presented in transparent plastic jars served with individual wooden tasting sticks. Respondents tasted as much honey as they considered necessary for its evaluation. During the tasting stage, respondents were allowed to communicate only with moderators.

#### 3.1.3. Questionnaire

The survey filled out at the product tasting stage contained three parts.

The first part of the questionnaire included a list of various sensory characteristics of honey, describing its taste, aroma, and textural properties. Respondents evaluated the degree of manifestation of the characteristics in the tasting samples on a scale from 0 to 6, where 0 — complete inconsistency of the taste, aroma or texture characteristic of honey, 6 — maximum manifestation of the characteristics in the sample. For example, if respondents felt that the honey exhibited a very strong sweetness compared to other samples, they could give the sample a score of 6 out of 6 on the sweetness scale. In total, the following 21 characteristics were considered:

1. Colour intensity;
2. Odour intensity;
3. Taste intensity;
4. Presence of foreign flavours;
5. Crystallization;
6. Florality;
7. Fruitiness;
8. Berryness;
9. Herbivory;
10. Woodiness;
11. Spiciness;
12. Tangibility of the taste or odour of wax in honey;
13. The sensation of artificial sugar;
14. Sweetness;
15. Bitterness;
16. Sourness;
17. “Peppercorn”, the tickling sensation in the throat after the honey tasting;
18. Duration of the aftertaste;
19. “Honey plant”, the accordance between the expected sensation of honey and the actual taste of the main component;
20. Tartness;
21. The feeling of astringency.

In the second part of the questionnaire, respondents rated the perceived taste of honey samples on a scale from 0 to 6, and the perceived quality of honey on a scale from 1 to 7. Participants could also leave their comments on their scores in text format.

#### 3.1.4. Data analysis

To identify the interdependence of the sensory characteristics of honey, a Spearman correlation matrix was constructed to capture the correlations between the variables. To combine the 21 sensory characteristics into larger structures, we conducted an exploratory factor analysis (EFA). The term EFA refers to two distinct models with different objectives and computational methodologies, specifically, principal component analysis (PCA) and common factor analysis (Watkins, 2018). In the present study, common factor analysis was chosen for several reasons. Firstly, it offers enhanced capabilities for identifying hidden relationships within a data set, unlike PCA (see, for example, Gorsuch, 1990; Widaman, 1993). Additionally, this analytical approach has previously demonstrated its effectiveness in food preference studies (Sautron et al., 2015). Exploratory factor analysis, using parallel analysis with a Varimax rotation, was utilized for investigating underlying constructs of sensory characteristics that respondents were asked about during the questionnaire. Factors with eigenvalues above one and above the corresponding eigenvalues of the parallel analysis were accepted. The exclusion criteria for items and factors were as follows: items with a factor loading value below 0.4 were excluded. Items that load on more than one factor, as well as items that are not loading on a factor were considered redundant, and thus, were removed. To evaluate the coherence of the items within each factor, internal consistency was tested with Cronbach’s alpha scores (see, for example, DeVellis, 2017). The Cronbach’s Alpha values were interpreted in the following manner: α ≥ 0.9 “Excellent”, 0.7 ≤ α < 0.9 “Good”, 0.6 ≤ α < 0.7 “Acceptable”, 0.5 ≤ α < 0.6 “Poor”, and α < 0.5 was categorised “Unacceptable”.

##### Regression analysis

To examine the relationship between the factors, revealed by the factor analysis, with WTP, the perceived honey quality and taste, the following (1)-(3) multiple linear regressions were used:

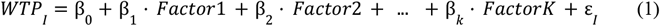

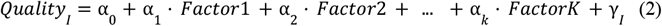

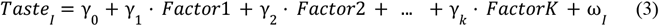

where:

*WTP*_*I*_— WTP for the subject *i* for the honey sample *m*;

*Quality*_*I*_— quality of the honey sample *m*, perceived by subject *i*;

*Taste*_*I*_— taste of the honey sample *m*, perceived by subject *i*;

*Factor1*,*…, FactorK* — factors formed from sensory characteristics of honey using factor analysis;

ε _*I*_, γ _*I*_, ω_*I*_— idiosyncratic shocks to each subject *i* for the honey sample *m*.

All data analysis was executed in R Studio, version 1.1.456.

### 3.2. Neuromarketing experiment

#### 3.2.1. Participants

The second stage of the study involved 25 healthy volunteers (9 men and 16 women) aged 21 to 55 years (mean age 28.2 ± 7.9 years) without contraindications to the study. This group of participants did not overlap with the group from the behavioural experiment. All participants were recruited for the experiments through social media invitations and snowball sampling. Eligibility criteria included the absence of food allergies or other medical contraindications, as well as regular (at least once a month) consumption of honey. In order to minimize potential bias in WTP assessments due to differences in hunger levels (Briz et al., 2015), we asked each participant to rate their level of hunger on a 5-point Likert scale before the start of the experiment. Consequently, the results of this survey showed that no participants were hungry before the beginning of the experimental session. Additionally, we requested that all participants refrain from consuming coffee, tea, tobacco, and alcohol for at least 4 hours prior to the experiment to prevent any possible distortions in sensory perception and recorded EEG signals (see, for instance, Ehlers et al., 1989). The experimental procedure was approved by the local ethics committee, and every participant signed an informed consent form prior to the commencement of the experimental session.

#### 3.2.2. Stimuli

The same seven brands of flower honey that were tested in the behavioural experiments were tested in the neuromarketing experiments. The samples differed in price, but were identical in all other characteristics (honey type, year of collection, colour, etc.).

#### 3.2.3. Experiments procedure

The experimental session was conducted as follows: respondents were comfortably seated in front of a monitor, on which all necessary instructions and questions were displayed. Each participant in the experiment was asked to blindly taste 7 honey samples, which were given to the participants in a random order. The amount of each sample was approximately 20 g, which is sufficient to evaluate the taste and does not tire the respondent’s taste buds. After tasting each sample, the participants were asked to answer the following questions: 1) how much are they willing to pay for a 250 g jar of such honey in a store 2) how tasty (on a scale from 1 to 5) did they find the honey they had just tasted 3) how high (on a scale of 1 to 5) do they think the quality of the sample is, as well as the questions about the manifestation of the sensory characteristics that we have identified as factors in the behavioural part of the study. Between tasting different samples, participants were allowed to drink any amount of still water.

#### 3.2.4. EEG Recording

Throughout the experiment, participants were seated in front of a computer screen (1920 × 1200 ppi) placed approximately 60 cm away. They were instructed to respond to each sample using a keyboard. All instructions and participant responses were managed using a custom script developed by the authors, which utilized Yad, a command-line tool for creating graphical user interface (GUI) dialogues from shell scripts. This script is available upon request. EEG signals were continuously recorded using the portable wireless EEG system “NeuroPolygraph.” The system features 21 channels arranged according to the standard 10-20 electrode placement, covering positions FP1, FPZ, FP2, F3, FZ, F4, F7, F8, C3, CZ, C4, P3, PZ, P4, T3, T4, T5, T6, O1, OZ, and O2. Nylon cap was used to position the electrodes on the participants’ heads. To enhance signal conductivity, a specialized conductive gel was applied to the electrodes after placement, ensuring impedance remained below 20 kΩ. The EEG signals were sampled at 500 Hz, with all other parameters configured according to the manufacturer’s guidelines.

#### 3.2.5. Data analysis

##### 3.2.5.1. EEG processing

The data collected from all experiments were exported in EDF format and analysed using a custom Python script that utilized the MNE (Gramfort et al., 2013) and SciPy libraries (Virtanen et al., 2020). The offline processing pipeline included filtering, artifact rejection and segmentation to prepare the data for further analysis. Each participant’s EEG data were filtered using a bandpass filter with a lower cutoff frequency of 1 Hz and an upper cutoff frequency of 50 Hz. This filtering range was chosen to eliminate low-frequency noise, such as drift and movement artefacts, while retaining the brain activity signals typically observed within this frequency band.

To ensure data quality, all EEG segments were manually reviewed for artefacts. This involved visual inspection of the recordings to identify and remove artefacts caused by factors such as participant movement, muscle activity (e.g., swallowing), or eye blinks. This thorough manual review process allowed the researchers to ensure that only clean, artifact-free segments were included in the analysis, thereby enhancing the reliability and validity of the findings by minimizing the impact of non-neural signals.

For further analysis, 5-second segments were extracted from the EEG data. These segments corresponded to the periods when participants were contemplating their WTP for the honey sample after tasting it. As a result, seven segments were obtained per participant. The data from all segments and channels were then filtered in the alpha (8-12 Hz) and beta (13-30 Hz) frequency ranges using the fast Fourier transform method. Subsequently, the total signal power for each electrode was calculated in both frequency ranges.

The processed data from each electrode were referred to as neurometrics (Semenova et al., 2023). For the subsequent WTP analysis, the neurometric showing the highest statistically significant correlation with WTP was selected.

##### 3.2.5.2. Modelling

The influence of factors on consumers’ WTPfor honey was studied using regression analysis. It was decided to conduct a research using a two-stage linear regression. First, we used the declared perceived quality of the tasted honey as a dependent variable, and the explanatory variables were the metrics of the sensory characteristics that respondents were asked about after tasting each sample, in particular: intensity of sweet taste, intensity of fruity and berry taste, intensity of sour taste, tartness, absence of foreign odours and components, as well as sensation of the heterogeneous structure of honey. The general form of the estimated model is presented in equation (4):

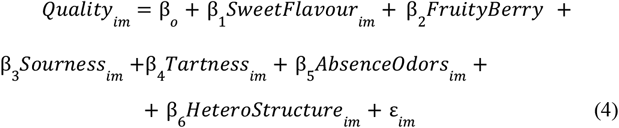

where:

*Quality*_*im*_ – the quality of honey sample *m* declared by individual *i*;

*SweetFlavour* _*im*_ – intensity of sweet taste (1 - not felt at all, 5 - felt very strongly) of a honey sample *m*, declared by the individual *i*.

*FruityBerry*_*im*_– intensity of fruity and berry flavour (1 - not felt at all, 5 - felt very strongly) of a honey sample *m*, declared by the individual *i*.

*Sourness*_*im*_– intensity of sourness (1 - not felt at all, 5 - felt very strongly) of a honey sample *m*, declared by the individual *i*.

*Tartness*_*im*_– intensity of tartness (1 - not felt at all, 5 - felt very strongly) of a honey sample *m*, declared by the individual *i*.

*AbsenseOdors*_*im*_– absence of foreign odours and components (1 - not felt at all, 5 - felt very strongly) of a honey sample *m*, declared by the individual *i*.

*HetroStructure*_*im*_ – sensation of the heterogeneous structure of honey (1 - not felt at all, 5 - felt very strongly) of a honey sample *m*, declared by the individual *i*.

ε_*im*_ – an idiosyncratic shock to each subject i for the honey sample m;

Next, a model for assessing consumers’ WTP was constructed (5), where the consumer’s stated WTP for a 250-gram jar of honey was used as a dependent variable, and some personal characteristics of the individual (in particular, gender and age), the individual’s stated assessment of the taste of the honey sample, and the value of the quality of the honey sample estimated from equation (4) through sensory characteristics were included in the model. In addition, neurometrics F4 in the beta range was included in the model as one of the dependent variables:

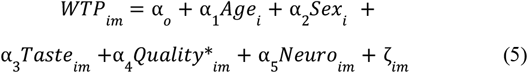

where:

*WTP*_*im*_– the stated WTP by individual *i* for a sample of honey *m*;

*Age*_*i*_– age of the subject i (full years);

*Sex*_*i*_ – dummy variable, indicating sex of the subject i, where 1— a male, 0 — a female;

*Taste*_*im*_– the taste of honey sample *m* declared by individual *i*;

*Quality*\*_*im*_– the fitted values of the variable quality in equation (4);

*Neuro*_*im*_– power of the EEG signal from the selected channel (F4 in beta range) for the subject i during the tasting of the honey sample m;

*ζ*_*im*_– an idiosyncratic shock to each subject i for the honey sample m;

## 4. Results

### 4.1. Results of behavioural experiments

The results demonstrate significant correlations between the sensory characteristics of honey and the dependent variables: the perceived quality of the product and respondents’ WTP for it. The accordance between the expected perception of honey and the actual taste of the main component, the intensity of flavour, the duration of aftertaste, and the tickling sensation in the throat after the honey tasting have a positive effect on the perceived quality of honey, while the sensation of extraneous flavours and added sugar shows the opposite trend. Interestingly, neither “Crystallisation” (r_s_=-0.04, p-value=0.60) nor degree of “Sweetness” (r_s_=0.01, p-value=0.85) statistically significantly influenced consumers’ perception of honey samples. The correlation matrix reveals the presence of relationships between different sensory characteristics of honey: e.g. “Fruitiness” and “Berryness” (r_s_ = 0.59, p-value<0.001), “Woodiness” and “Herbivory” (r_s_ = 0.44, p-value<0.001), “Bitterness” and “Peppery” (r_s_ = 0.57, p-value<0.001) etc. Thus, according to the results of the correlation analysis, it seems possible to combine the considered sensory characteristics into several groups. Grouping of the components that fully describe the flavour portrait of honey will significantly reduce the number of variables, which is required to conduct time-consuming and labour-intensive study of honey perception.

### 4.2. Results of the factor analysis

Parallel analysis on the sensory characteristics resulted in a reduced 16-item scale, with 6 factors accounting for a total of 47% of variance (see Table 1 for more details on the results of the exploratory factor analysis). Based on the criteria set out for item and factor reduction, five items (“Colour intensity”, “Odour intensity”, “Florality”, “Aftertaste” and “Herbivory”) were removed due to cross-loading on two or more factors.

**Table 1.** Results of exploratory factor analysis, showing the rotated component matrix with sensory characteristics loadings on six factors.

| Items | Factors (Exp.Var. of Data, "=Cronbach's Alpha) |  |  |  |  |  |
| --- | --- | --- | --- | --- | --- | --- |
|  | Factor 1 (15%,<br>=0.81) | Factor 2<br>(8%,<br>=0.55) | Factor 3<br>(8%,<br>=0.55) | Factor 4<br>(7%,<br>=0.7) | Factor 5<br>(5%,<br>=0.67) | Factor 6<br>(4%,<br>=-) |
| Bitterness | 0.74 |  |  |  |  |  |
| Tartness | 0.69 |  |  |  |  |  |
| Peppercorn | 0.65 |  |  |  |  |  |
| Spice | 0.63 |  |  |  |  |  |
| Woodiness | 0.49 |  |  |  |  |  |
| Artificial sugar |  | -0.65 |  |  |  |  |
| Foreign tastes |  | -0.62 |  |  |  |  |
| Taste intensity |  |  | 0.68 |  |  |  |
| Sweetness |  |  | 0.58 |  |  |  |
| Fruitiness |  |  |  | 0.75 |  |  |
| Berry |  |  |  | 0.74 |  |  |
| Wax |  |  |  |  | 0.61 |  |
| Astringency |  |  |  |  | 0.49 |  |
| Sourness |  |  |  |  |  | 0.53 |

The characteristics loading on Factor 1 represented variables related to the tartness of honey, thus, we named Factor 1 “Tartness”. Factor 2 included characteristics related to the content of foreign components in honey. Since the sign of the loadings was negative, Factor 2 was named “Absence of extraneous odours and components”. As for Factor 3, it consisted of such honey characteristics, as “Taste intensity” and “Sweetness”, and therefore, it was named “Intensity of sweet flavour”. The fourth factor consisted of characteristics representing fruity or berry shades of honey taste, therefore, this factor was called “Fruitiness”. Factor 5 was named “Sensation of the heterogeneous structure of honey”, because it included the characteristics of wax and astringency. Lastly, we named Factor 6 “Sourness”, because it included the only characteristic — “Sourness”. The overall reliability of the 16-item scale was good (Cronbach’s Alpha = 0.722). Likewise, the internal consistency reliability of each of the six factors was acceptable to good.

### 4.3 Results of linear regressions estimations

Table 2 lists the OLS-estimates for the equations (1)-(3). As expected, the factors formed on the basis of sensory characteristics best describe the perceived quality of honey, showing for this model the highest R^2^ value of 0.668. Next comes the model with the perceived taste as the explained variable with R^2^ of 0.454%. Finally, in the model where WTP was the explained variable, the proportion of explained variance was 24%.

**Table 2.** OLS-estimates for factors, influencing WTP, quality and perceived taste of the honey on the data of behavioural experiments.

| Factors | WTP<br>(1) | Quality<br>(2) | Perceived<br>taste<br>(3) |
| --- | --- | --- | --- |
| Tartness | 24.293**<br>(11.815) | 0.548***<br>(0.090) | 0.448***<br>(0.117) |
| Absence of extraneous odours<br>and components | 37.705***<br>(12.014) | 0.317***<br>(0.091) | 0.345***<br>(0.119) |
| Intensity of sweet flavour | 51.002***<br>(12.496) | 1.293***<br>(0.095) | 1.113***<br>(0.124) |
| Fruity and berry flavour | 28.559**<br>(12.450) | 0.496***<br>(0.095) | 0.448***<br>(0.123) |
| Sensation of the heterogeneous<br>structure of honey | 29.556**<br>(12.903) | 0.126<br>(0.098) | 0.148<br>(0.128) |
| Sourness | -13.990<br>(13.704) | -0.118<br>(0.104) | -0.044<br>(0.136) |
| Constant | 205.031***<br>(10.379) | 4.484***<br>(0.079) | 3.629***<br>(0.103) |
| $R^2$ | 0.240 | 0.668 | 0.474 |
| Adjusted $R^2$ | 0.210 | 0.655 | 0.454 |
| N | 159 | 159 | 159 |

*Note: significance: \*\*\* = $p < 0.01$ , \*\* = $p < 0.05$ , \* = $p < 0.1$ . Standard errors in parenthesis*

Estimation of the equation (4), in which we explained the perceived quality of honey through aggregated sensory characteristics, gave the following results: the intensity of fruity and berry taste, intensity of sweet taste, tartness and absence of foreign odours have a statistically significant positive effect on the perceived quality (Table 5). In general, the aggregated sensory characteristics explained about 50% of the variation in the perceived quality of honey. It is also worth noting that the results of assessing the perceived quality of honey using neuromarketing experiments, in general, are consistent with the results of assessing the perceived quality of honey using behavioural experiment data, where the same regressors had a statistically significant effect on the dependent variable: the intensity of fruity taste, intensity of sweet taste, tartness, and foreign odours.

**Table 5.** OLS-estimates for factors, influencing quality of the honey on the data of neuromarketing experiments.

|  | Quality |
| --- | --- |
| Intensity of sweet flavour | 0.129*<br>(0.077) |
| Fruity and berry flavour | 0.226***<br>(0.057) |
| Sourness | -0.084<br>(0.074) |
| Tartness | 0.153***<br>(0.058) |
| Absence of extraneous odours and components | 0.465***<br>(0.052) |
| Sensation of the heterogeneous structure of honey | -0.072<br>(0.050) |
| Constant | 3.214***<br>(0.366) |
| R<sup>2</sup> | 0.509 |
| N | 175 |

*Note: significance: \*\*\* = $p < 0.01$ , \*\* = $p < 0.05$ , \* = $p < 0.1$ . Standard errors in parenthesis*

Next, having obtained the estimated value of the perceived quality of honey through the aggregated sensory characteristics, we used it as one of the explanatory variables in the WTP assessment model (5). This step allowed us to avoid the problem of multicollinearity due to the high correlation between the variables of perceived quality and perceived taste (the correlation coefficient between these variables was 0.81, while the correlation coefficient between perceived taste and the estimated value of perceived quality in equation (4) (variable Quality*) was 0.69).

The results of the model (5) estimation are presented in Table 6. The male gender, the perceived taste, and the perceived quality, which includes the assessments of the sensory characteristics of honey, have a statistically significant positive effect on the willingness of consumers to pay for honey, while the increase in the activity of the frontal zone of the brain, expressed through the F4 neurometrics in the beta range, has a statistically significant negative effect on the willingness of consumers to pay. In particular, on average, all other factors being equal, men are willing to pay 33.5 monetary units more for honey; each additional point of the subjective assessment of taste increases the WTP by 48.7 monetary units; an increase in the value of the perceived quality leads to an increase in the WTP by 31.7 monetary units; finally, an increase in the activity of the frontal zone of the brain reduces the willingness of consumers to pay for honey.

**Table 6.** OLS-estimates for factors, influencing WTP for honey, on the data of neuromarketing experiments.

|  | WTP |
| --- | --- |
| Age | -0.180<br>(0.823) |
| Sex | 33.493**<br>(14.255) |
| Taste | 48.735***<br>(6.885) |
| Quality* | 31.704***<br>(9.856) |
| Neuro (F4 beta) | -377.788**<br>(186.085) |
| Constant | -65.435**<br>(31.870) |
| Adjusted R<sup>2</sup> | 0.506 |
| N | 175 |

*Note: significance: \*\*\* = $p < 0.01$ , \*\* = $p < 0.05$ , \* = $p < 0.1$ . Standard errors in parenthesis*

## 5. Discussion

This study presents a multi-stage approach to assess the influence of honey sensory characteristics on consumers’ WTP for it. The first step in assessing the perceived quality of honey was to conduct a series of behavioral experiments in which respondents blindly tasted honey samples and assessed their 21 sensory characteristics. Next, taking into account the complexity of the task of identifying individual taste characteristics by ordinary consumers, we conducted exploratory factor analysis and combined 21 sensory characteristics of honey into 6 factors (“Astringency”, “Absence of foreign odors and components”, “Intensity of sweet taste”, “Fruity”, “Sensation of heterogeneous structure of honey”, “Acidity”). Given the high proportion of explained variance, we concluded that six aggregated characteristics obtained as a result of factor analysis are sufficient to form a complete taste portrait of honey.

Then, using the characteristics obtained from the factor analysis results, we estimated linear regression models of the dependence of willingness to pay, perceived taste and quality on the characteristics described above. Having estimated such models, we found that the aggregated sensory characteristics best describe the perceived quality of honey. We used this information further in the neuromarketing part of the study.

In the second, neuromarketing part of the study, we conducted a series of laboratory experiments using EEG, during which respondents were also asked to taste honey blindly and assess their own willingness to pay, the perceived taste and quality of honey, and a set of 6 sensory characteristics that we obtained through factor analysis based on the behavioral experiments described earlier. The data obtained as a result of the neuromarketing experiments were used to construct a two-stage linear regression, where in the first step the perceived quality was used as a dependent variable, and the 6 sensory characteristics were used as regressors, and in the second step the assessed perceived quality was one of the explanatory variables for consumers’ willingness to pay for honey. Notably, the findings about sensory characteristics from neuromarketing experiments aligned closely with those from behavioural experiments, reinforcing the robustness of the results. Both methods identified the same regressors — fruity taste, sweet taste, astringency, and foreign odours — as statistically significant predictors of perceived quality. This consistency across methodologies underscores the reliability of sensory attributes as determinants of honey quality. Investigating the neural mechanisms underlying these sensory perceptions could provide deeper insights into how consumers process and evaluate sensory information. For instance, are certain tastes or odours processed more quickly or intensely in the brain, and how does this affect overall preference? Moreover, cross-cultural studies could be conducted to determine whether these findings hold across different consumer groups with varying taste preferences and cultural backgrounds.

The results from the regression model for WTP reveal that men exhibit a higher WTP for honey compared to women, suggesting potential gender-based variations in consumer behaviour. This aligns with existing research highlighting gender as a key demographic factor in purchasing decisions and underscores the need for further exploration of how gender influences product valuation. Understanding these differences could help tailor marketing strategies to better address the preferences of different consumer segments.

The study also highlights the critical role of sensory attributes, particularly perceived taste and quality, in driving WTP. Consumers place a high value on the sensory experience of honey, which directly influences their WTP a premium. For producers, this emphasizes the importance of prioritizing taste and overall quality as a strategic approach to justifying higher prices and differentiating their products in competitive markets. Enhancing sensory attributes could be achieved through careful selection of floral sources, optimized processing techniques, and rigorous quality control measures.

A particularly intriguing finding is the negative relationship between increased activity in the frontal zone of the brain (measured through F4 neurometrics in the beta range) and WTP. This suggests that heightened cognitive engagement during product evaluation may lead to more critical or cautious decision-making, ultimately reducing the WTP. This neurophysiological insight adds a new dimension to our understanding of consumer behaviour, demonstrating that neural activity can serve as a predictor of purchasing decisions. It also opens up opportunities for the application of neuromarketing techniques, where insights into cognitive processes could be leveraged to design more effective marketing campaigns.

The integration of behavioural and neurophysiological data in this study represents a significant methodological advancement, providing a more comprehensive understanding of consumer decision-making. By combining traditional survey-based approaches with EEG measurements, this research demonstrates the value of interdisciplinary methods in capturing the complex interplay between sensory perceptions, cognitive processes, and economic behaviour. This approach could be extended to other food products or consumer goods to further validate its applicability.

In conclusion, this study highlights the multifaceted nature of consumer decision-making, demonstrating that both sensory and neurophysiological factors play a significant role in shaping WTP. By integrating these insights, producers and marketers can develop more effective strategies to meet consumer preferences and enhance market competitiveness.

## 6. Conclusion

The findings provide new knowledge of the honey choice behaviour, which can contribute to the better understanding of the honey perception by consumers and support food researchers and the food industry. Underlying the sensory characteristics that positively influence consumer perception of a product is crucial for solving various challenges in honey marketing. This knowledge can be applied to effectively design honey positioning, development of an impactful communication strategy, and creation of attractive packaging that enhances the product’s appeal to potential customers.

Reducing the number of analysed characteristics of honey allows us to simplify the task for respondents who are not professional tasters, identifying the subtlest differences in characteristics, but simple consumers with specific requests for a certain quality of the product. This solution contributes to the reduction of the experiment duration, because the respondents have to evaluate only 6 characteristics instead of 21. This fact will be potentially significant in the context of conducting neuromarketing research, where the respondents are forced to stay in specific equipment for a long time.

Further investigations of consumers’ perception of honey should be carried out to get a deeper insight into the perceived quality of the product, as a metric explained by a subjective assessment of a set of sensory characteristics and influencing consumers’ WTP.

